# NIR Phosphorescent Oxygen Sensors in Natural Hydrogel Matrices

**DOI:** 10.1101/2025.05.17.654688

**Authors:** Waqas Saleem, Ananthakrishnan Soundaram Jeevarathinam, Rebekah Lindblade, Michael J. McShane

## Abstract

Understanding the effects of sterilization and other treatments over different hydrogels used as matrices for oxygen-sensitive microparticles is essential in designing biocompatible phosphorescent metabolite sensors. In this study, we dispersed oxygen-sensitive microparticles in hydrogel matrices from various natural sources, such as gelatin, alginate, collagen, and Albumin. Subsequently, we comprehensively analyzed their mechanical and rheological properties and oxygen sensitivity before and after treatments, including autoclave and E-beam sterilization and exposure to serum and cell culture conditions. Our findings, encompassing oxygen sensitivity, compression testing, and rheological profiling, consistently indicate that the E-beam sterilization method yields the most reliable results when applied to alginate and Albumin (BSA) matrices containing oxygen-sensing microparticles. Furthermore, BSA gels exhibited robust stability and performance characteristics, demonstrating minimal structural alterations following post-treatment conditions.

## Introduction

Hydrogel materials, characterized by their water-absorbing and retaining properties, owe their generally excellent biocompatibility to a hydrophilic nature like the extracellular matrix (ECM) of biological tissue. Researchers have categorized hydrogels based on various parameters, including their origin and physical and chemical attributes. Origin-wise, hydrogels fall into three primary categories: synthetic polymers (e.g., polyethylene glycol, polyacrylic acid, polystyrene sulfonate), natural polymers (e.g., alginate[1], [2], [3], [4], [5], [6], [7], [8], [9], [10], hyaluronic acid[11], carrageenan[12], [13], starch[14], [15], [16], gelatin[2], [17], serum albumin[18], [19], collagen[2], [4], DNA-based hydrogels[20], and hybrid polymers[21]. Hybrid hydrogels can result from grafting synthetic groups onto natural polymers. They can also emerge from combining non-organic components (e.g., metal or clay) with hydrophilic polymers, such as starch and silver nanoparticles. [16]

Researchers have characterized hydrogels as biological (natural/biomaterial), synthetic, or hybrid based on their composition and derivations. [22] Biological hydrogels are polymers derived from natural sources like alginate, agarose, collagen, carrageenan, chitosan, fibrin, and hyaluronic acid (HA). These naturally occurring polymers offer inherent biocompatibility and biodegradability but may lack precise control over mechanical and structural properties. In contrast, synthetic hydrogels are formed by the chemical synthesis of polymeric networks, such as polyethylene glycol (PEG), polyhydroxyethyl methacrylate (pHEMA), and polyvinyl alcohol (PVA). These synthetic materials offer greater tailorable control over molecular structure, crosslinking density, biodegradation, and mechanical strength. Biohybrid hydrogels bridge the gap between these two categories by integrating an active biological entity (naturally derived peptide, genetically engineered protein, enzyme, or growth factor) into synthetic networks. This synergistic approach allows for retaining the inherent biocompatibility of natural polymers while also enabling precise control over biological properties like high affinity and specific binding, as well as tailored mechanical strength and stimuli-responsive behavior. Examples include PEG and PVA hydrogels modified with amino acid sequences (RGD/RGDS peptide), which enhanced cellular adhesion to these hydrogels.

Natural hydrogels have garnered significant attention in biomedical research due to their inherent biocompatibility and biodegradability. They form an environment reminiscent of natural soft tissue ECM, creating a conducive milieu for cellular activities. Consequently, natural hydrogels have found extensive utility in various applications, including tissue engineering [23], [24], [25], drug delivery[26], [27], [28], [29], [30], and vaccine[31], [32], [33], [34], [35] development. Their versatility allows them to be scaffolding materials for encapsulating cells, drugs, cellular products, and optically responsive biosensors.

Among the natural polymers derived from polysaccharides, alginate is a prominent choice. This anionic polymer, extracted from brown seaweed, has undergone extensive exploration for biomedical and pharmaceutical applications. Alginate is extensively used as a gelling agent due to its biocompatibility, cost-effectiveness, low toxicity, and rapid gelation properties. Alginate gelation occurs only upon its chemical interaction with divalent ions (Ca^2+^, Br^2+^, Sr^2+^) or trivalent ions (Fe^3+^ and Al^3+^). [36] The negatively charged carboxyl groups of alginate polymer chains attract the positively charged calcium ions, resulting in an overall "egg-box structure" where the egg represents calcium ions while the wavy box represents adjacent anionic polymer chains of alginate. [36], [37] However, the replacement and subsequent release of divalent cations from the crosslinked hydrogel structure due to ion exchange can weaken alginate gels over time when immersed in media containing high concentrations of non-gelling ions (Na^+^ and Mg^2+^), promoting gel instability. [36], [38]Conversely, monovalent complex ions such as phosphate and citrate can act as a potent complex-forming agent, sequestering Ca^2+^ ions and disrupting gelation. [36] Such limitations render ionically-crosslinked alginate hydrogels prone to inconsistent physical properties and accelerated degradation. [39], [40]

Researchers have pursued diverse strategies for enhancing their properties in response to the limitations of ionically crosslinked alginate gels. Initial efforts were focused on modifying alginate side chains. This includes the introduction of methacrylates for photo-crosslinking [41] and incorporating tetrazine and norbornene groups to enable click-chemistry-based covalent crosslinking. [42] Similarly, another research group reported carbodiimide chemistry to crosslink alginate and gelatin polymers covalently43] while other researchers were interested in designing a cell-laden polymer mixture of alginate-gelatin-collagen for controlled degradation. [2] Similarly, ionically crosslinked double network hydrogels of alginates with k-carrageenan [44] or polyacrylamides [45] were also reported. These approaches have yielded stable hydrogel formulations with more consistent and, in some cases, tunable mechanical properties tailored to specific applications.

Conversely, protein-based natural hydrogels derived from biopolymers like gelatin, bovine serum albumin (BSA), and collagen offer distinct advantages, including enhanced yield strength, compatibility with soft tissues, and the ability to more closely mimic ECM microarchitecture. [2], [19], [46] Albumin, which accounts for over 50% of blood plasma proteins, is an economical source of water-soluble proteins. Albumin hydrogels exhibit high mechanical stability, shape memory properties, and provide a xenofree platform for tissue engineering and drug delivery applications.[18], [47], [48], [49]

Hydrogels, characterized by their inherent high water content, exhibit exceptional optical transparency across a broad spectrum encompassing the visible and near-infrared regions. [50] This unique combination of optical clarity, biocompatibility, tunable mechanical properties, and gas permeability within their polymeric networks has positioned hydrogels beyond their traditional role in contact lenses [51], making them the material of choice for diverse applications. Examples include hydrogel-based photonic devices [52], [53] such as thin hydrogel films [54], [55], optical hydrogel fibers [56], and waveguides [57], [58]; plasmonic/photonic-hydrogel sensors [59], [60], [61] and implantable electrochemical-[62], [63], [64]/optical-based [65], [66], [67], [68], [69] biosensors.

We have published several examples of fully insertable biosensors which are optically transduced via phosphorescent-based lifetime readouts provided against target metabolites such as oxygen [1], [70], [71], glucose [8], [71], [72], lactate [67], [71], and uric acid [73] molecules. These sensors incorporate a naturally occurring, long-lifetime, oxygen-sensitive phosphor dye [Palladium (II) benzoporphyrin (PdBP)], which is loaded within alginate microparticles and subsequently dispersed in a hydrogel matrix. Long-lifetime phosphor dyes have delayed emission (in microseconds) that temporally separates them from the autofluorescence of native proteins (in nanoseconds), thereby minimizing the background interference from the skin. These biosensors are also fully insertable via a hypodermic needle, thanks to their grain-of-rice-sized footprint that offers minimal discomfort to the recipient and eliminates the need for a specialized caregiver for subcutaneous administration.

Realizing fully optimized and functional insertable biosensors necessitates addressing several critical challenges. These include but are not limited to ensuring biocompatibility of polymeric constituents [74], [75], sterilization impact and compatibility [76], design features facilitating needle-based delivery [77], [78], [79], [80], [81], and compatibility between material-tissue interface and mechanical properties [82], [83]. **Figure 1** depicts substantial variability in elastic modulus values reported using different measurement techniques across different skin regions, genders, and ages. [84] Similarly, Kalra [85] also provided a compiled list of elastic moduli from different skin layers, determined using diverse techniques. Notably, the elastic modulus values varied within the same body region and depended on the assessment technique. Moreover, the heterogeneity of results also highlights the limitations of a "one-size-fits-all" approach toward realizing an optimal biosensor. Consequently, a versatile and broader framework would be beneficial for settling different design criteria ranging from tunable mechanical properties to robust post-sterilization sensor performance.

**Figure 1:**
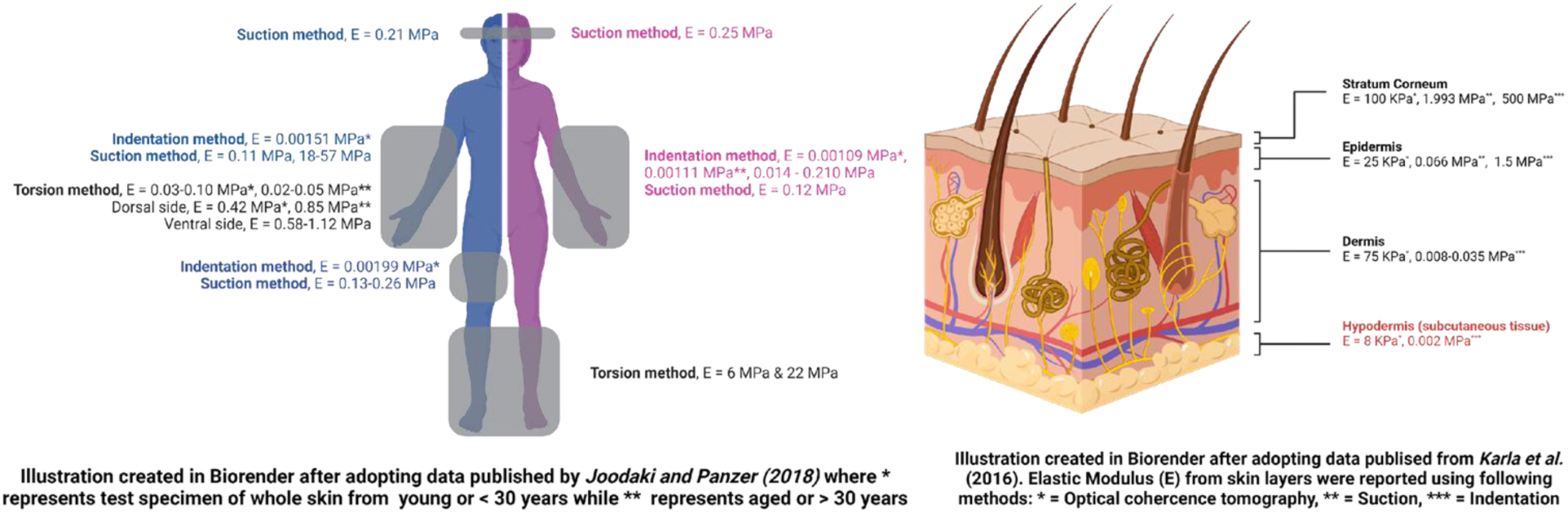
(LEFT) Graphical illustration of reported Elastic Modulus (E) of whole skin from male (blue) and female (purple), or both (black), located at different body regions and from different age groups - measured using different techniques. (RIGHT) Figurative representation of “E” reported in literature for different layers of the human skin.

This paper specifically focuses on using natural hydrogels from various sources to develop insertable optical-based oxygen sensors subjected to diverse sterilization methods and treatment conditions. Later, the sensors’ sensitivity responses and mechanical and rheological changes were determined and compared against each other, elucidating the effects of various sterilization methods and treatment conditions. Such comprehensive analyses are critical for determining 1) the best choice material of natural hydrogel with optimal oxygen sensitivity, 2) the robust sterilization method, and 3) the appropriate mechanical properties of insertable oxygen sensors for the targeted insertion site.

## Results and Discussion

As part of our goal of developing tissue-insertable optical oxygen sensors, we evaluated the potential of various natural hydrogel materials as matrix materials to carry the sensing chemistry. We explored using naturally-derived proteins such as gelatin, collagen, and Albumin by incorporating these biomaterials into the design of oxygen sensors embedded within different polymeric matrices: alginate, bovine serum albumin (BSA), and gelatin-alginate-collagen (GAC) gels. This diversification enabled us to expand our portfolio of materials usable for creating oxygen sensors with distinct physical characteristics and mechanical strength. **Figure 2** illustrates the overall strategies for developing and evaluating these oxygen sensors within naturally derived biopolymer matrices. A key consideration in this study was understanding the effects of different post-fabrication treatments on the chemical and physical properties of the materials, including the oxygen-sensing responses of the sensors.

**Figure 2:**
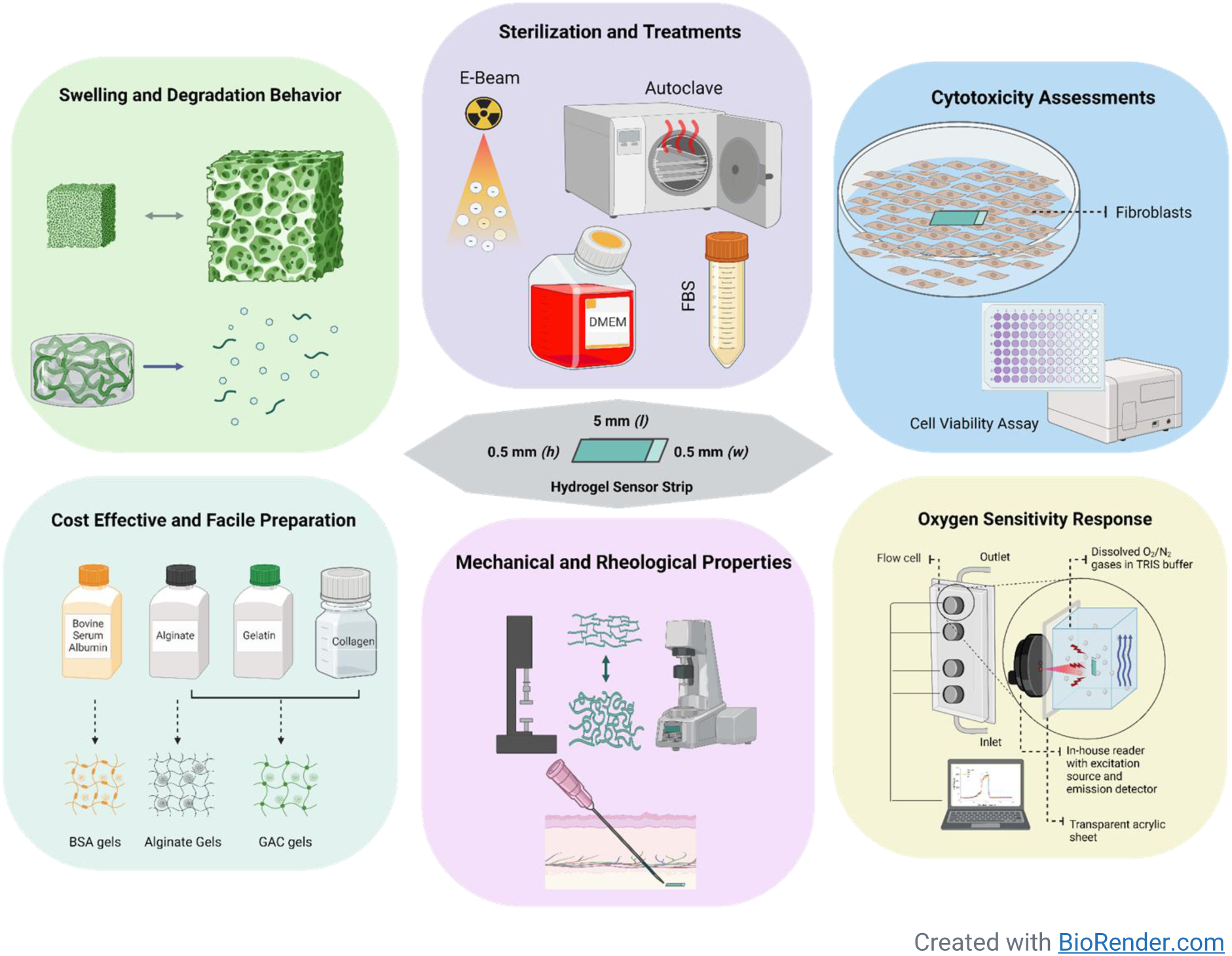
Schematic representation of the experimental design. Microparticles with encapsulated phosphor were dispersed in three different hydrogel matrices, which were then assessed for cytotoxicity and swelling as well as stability. Mechanical properties (compression and shear modulus) response to oxygen were measured and compared for native and samples exposed to sterilization or incubation in various media to elucidate the effects of treatments.

The fabrication process commenced with the synthesis of alginate microparticles containing oxygen-sensitive Pd (II) benzoporphyrin dye using a water-in-oil emulsion technique, a method that was developed in previous work. Subsequently, we applied nanofilm coatings comprising five successive bilayers of oppositely charged polyelectrolytes onto the microparticle surfaces to stabilize the alginate and retain the encapsulated phosphor. We evaluated the distribution of these microparticles’ sizes through the Nexcelom Cellometer, verifying the absence of aggregates formed during the microparticle fabrication process.

Next, we dispersed the oxygen-sensitive microparticles within the different natural hydrogel materials and crosslinked the matrices using methods appropriate to the materials involved. Depending on their intended use in various tests, the hydrogel matrices were crosslinked into monolithic slabs of varying thicknesses. Subsequently, circular discs were obtained to assess the physical responses of the gels, including swelling behavior, degradation profiles, and cytotoxicity. Furthermore, the sensors underwent different sterilization and other treatments, namely autoclaving, E-beam exposure, and immersion in various liquid media (DMEM cell culture medium, 10% fetal bovine serum, and 100% fetal bovine serum. We collected pre- and post-treatment data from each sensor within different matrices, evaluating oxygen sensitivity, mechanical properties, and rheological characteristics. Statistical analyses were conducted to discern the effects of various treatments within a given sample and across the samples.

### Physical Properties

#### 1. Physical characteristics of oxygen-sensing alginate microparticles

We employed Nexcelom’s mini-Cellometer device for analysis of size distribution and mean diameter calculation of oxygen-sensing alginate microparticles (MPs). **Figure 3** represents the microscopy and size distribution plot of MPs. The image revealed minimal microparticle aggregates in the sample, with a mean diameter of MPs around 9.2 microns.

**Figure 3:**
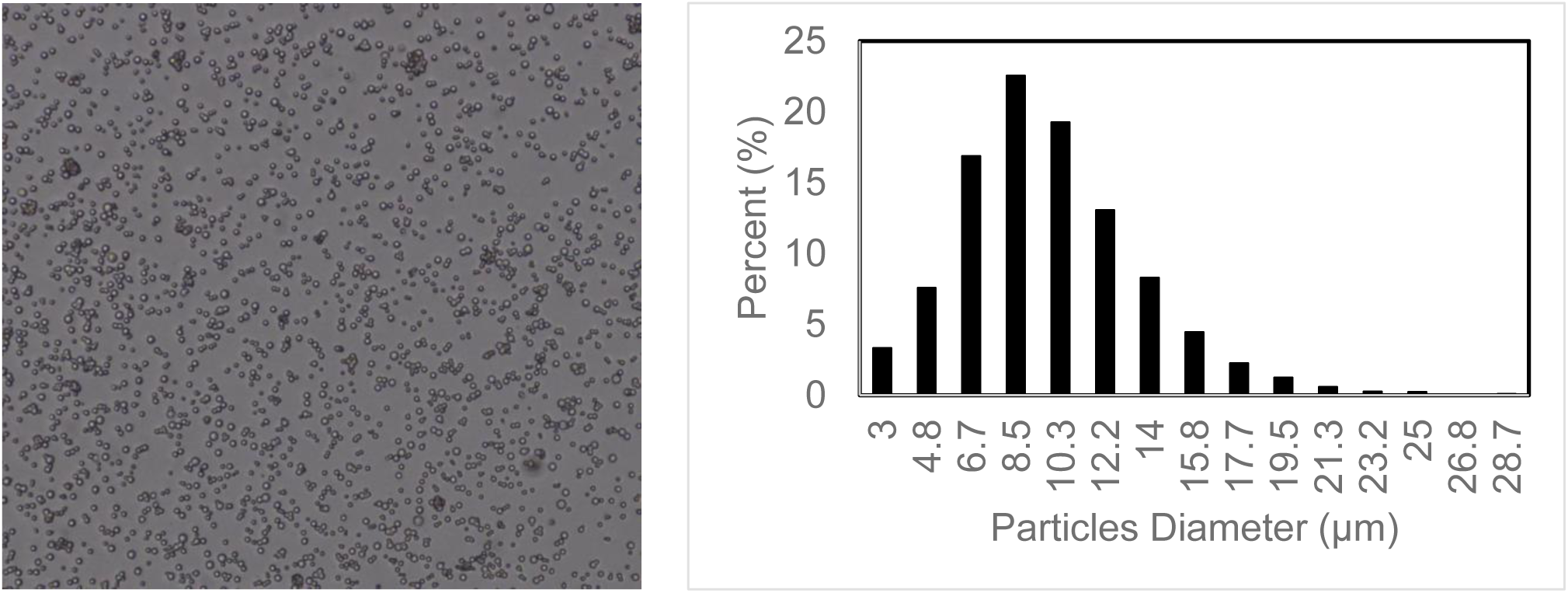
(LEFT) Microscopy of fabricated oxygen sensing alginate microparticles and (RIGHT) size distribution analysis of alginate microparticles.

#### 2. Swelling Behavior and Resilience of Hydrogels

We used scanning electron microscopy (SEM) to examine dehydrated hydrogel surface topology and microarchitecture, as depicted in **Figure 4**. Other research groups have separately reported SEM results from the exact composition of blank alginate [86] and BSA [49] gels, respectively, whereas a slightly different polymeric composition of blank GAC (4-1-1%) [87] gel was reported. No significant visual differences were observed between previous observations and our results. Interestingly, with the inclusion of oxygen-sensitive alginate microparticles in alginate and GAC hydrogels, significantly increased perforations were observed against its respective blank gel. In contrast, such differences were least prominent among the BSA gels. It is also critical to note here that SEM images do not precisely reflect the degree of porosity and microarchitecture of the native hydrated state of gels but provide researchers with additional insights to elucidate resulting structural changes/differences after varying polymeric contents and compositions.

**Figure 4:**
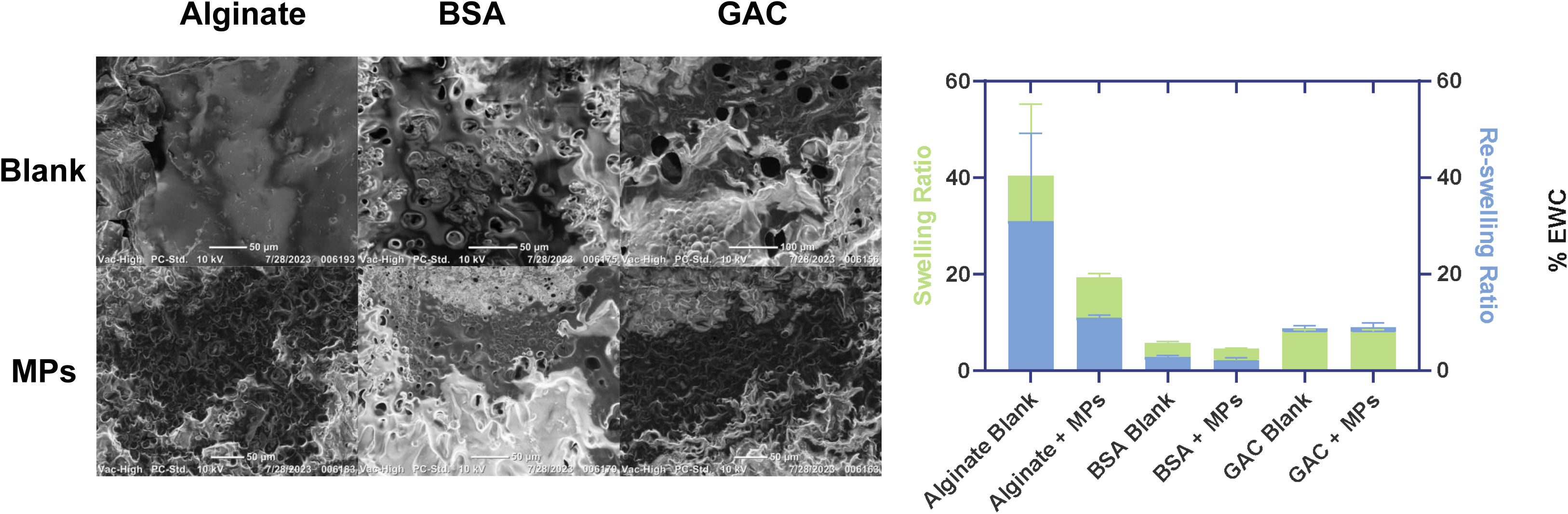
(LEFT) Scanning electron microscopy of dehydrated samples of blank (top row) and MP-containing hydrogels. (Middle) Swelling and re-swelling ratios of blank and microparticles–containing (MPs) hydrogels samples. (RIGHT) % Equilibrium Water Content (EWC) value of blank and MPs-containing hydrogels before and after freeze-drying of samples.

To evaluate the swelling behavior of these hydrogels, we subjected them to a swelling behavior assessment following freeze-drying by rehydrating the samples in their original storage buffer solution for at least 24 hours. Notably, blank- and MP-containing alginate gels exhibited higher swelling profiles and sample variations when compared with the other biomaterials tested. Previous work also found significant variability in the swelling behavior of Ca^2+^-crosslinked alginate gels when exposed to a similar concentration (10 mM) of CaCl_2_. Several factors, including the concentration of Ca^2+^ ions, the viscosity of the alginate solution, and the ratio of G-blocks to M-blocks in alginates, can influence the extent of swelling. [88] Intriguingly, the alginate gels displayed remarkable resilience in recovering their original water retention capacity. Based on % Equilibrium Water Content (EWC) values, blank alginate gels demonstrated only a 1.7% decrease, and MPs-containing alginate gels resulted in a 3.6% decrease after undergoing a single freeze-drying cycle.

In contrast, blank and MP-containing BSA gels exhibited the least swelling, with only partial recovery observed upon rehydration of the dehydrated samples. With a significant decrease in %EWC of 13% for blank BSA gels and a 16% decrease in MPs-containing BSA gels, this behavior can likely be attributed to protein denaturation and loss of its native conformation following the freeze-drying process.

Finally, the blank and MPs-containing GAC gels exhibited intermediate swelling profiles, with swelling and recovery falling between the alginate and BSA gels. Remarkably, these gels demonstrated nearly complete recovery of their swelling properties following rehydration of the dehydrated samples, with both blank and MPs-containing GAC gels only presenting a ∼1% increase in EWC values following the freeze-drying cycle.

#### 3. Degradation Profiles

Both blank alginate gels and MPs-containing alginate gels displayed limited stability when stored in aqueous media, as illustrated in **Figure 5**. This instability can be attributed to using ∼150 mM phosphate-buffered saline (PBS) buffer for hydrogel incubation. Inherently high concentration of sodium ions (130 mM NaCl) followed by potassium ions (2.7 mM of KCl) in PBS buffer would have resulted in the efflux of divalent Ca^2+^ ions responsible for the ionic crosslinking of adjacent polysaccharide polymer chains in the alginate matrix. Consequently, bulk erosion of the matrix occurred over a relatively short period in each case.

**Figure 5:**
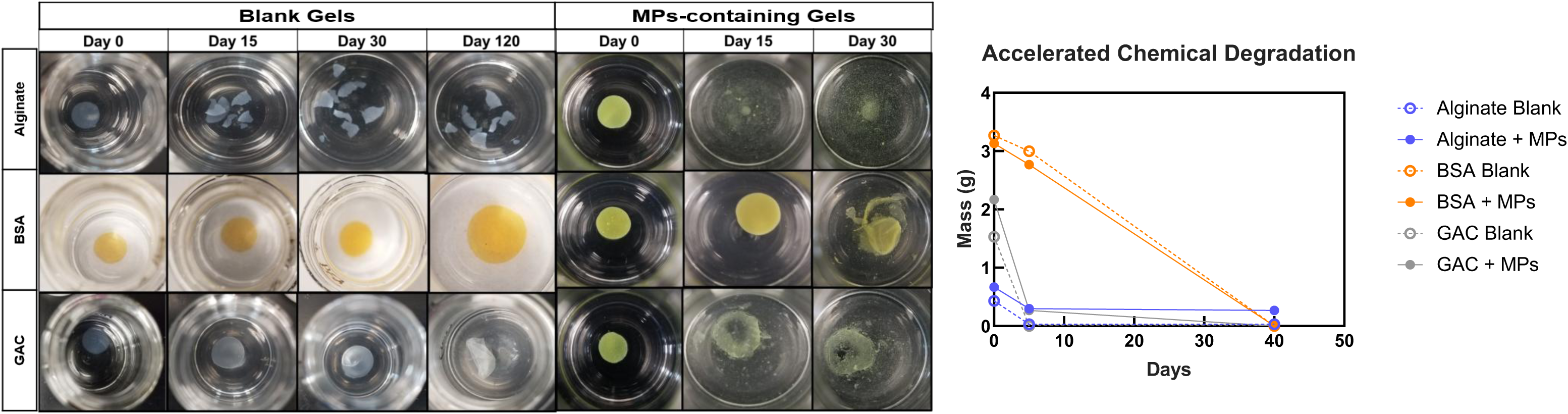
(LEFT) Stability testing of blank and MPs-containing hydrogels discs [5 (dia.) x 0.75 (h) mm] submerged in 5mL of 150 mM phosphate buffer saline (PBS) solution at room temperature was carried out. (RIGHT) Accelerated chemical degradation results from blank and MPs-containing hydrogel discs [3 (dia.) x 0.75 (h) mm] incubated in 0.1 M NaOH solution at 37°C for 40 days.

In contrast, blank GAC gels exhibited substantially better relative stability when immersed in PBS solution, with structural integrity maintained for at least 30 days. Subsequently, slight degradation became apparent around the 120-day mark. However, MP-containing GAC gels displayed substantially reduced stability compared to the native GAC gels, maintaining their structural integrity for approximately 15 days before exhibiting signs of bulk erosion. Weaker stability of MP-containing GAC gels can be attributed to 1) physical disruption of the continuous polymeric network in bulk gels due to microparticle addition [89], [90], 2) higher water content in microparticles than surrounding gel leading to dilution and reduction of matrix concentration [91], and 3) compositional changes affecting crosslinking density [92].

Interestingly, under PBS solution, blank BSA gels demonstrated the highest level of stability, resisting degradation for over 120 days, while MPs-containing BSA gels remained stable for about 15 days before experiencing bulk erosion. This enhanced stability in blank BSA gels can be attributed to the glutaraldehyde-based crosslinking method, which forms stable and robust crosslinks between primary amine functional groups. According to our hypothesis, the inclusion of alginate microparticles in BSA gels posed similar challenges as those experienced by MP-containing GAC gels where microparticles addition may have resulted in compositional change (affecting crosslinking density) and physical disruption of continuous polymeric chain network formations.

A similar trend was observed with accelerated chemical degradation studies; blank and MPs-containing BSA gels were most stable under hydrolytic conditions. On day 5, blank BSA gels reported an 8% loss in mass, while MPs-containing hydrogels reported a 12% decrease in mass. The second most stable hydrogel performance was demonstrated from MPs-containing alginate gels that reported a 55% reduction in mass, while blank alginate gels reported a 92% loss in mass. Interestingly, blank and MPs-containing GAC hydrogels reported 100% and 88% loss in mass, respectively. All gels were observed to be degraded entirely on day 40.

Based on these findings, we concluded that the crosslinked blank BSA gels exhibited superior hydrogel stability under physiological and hydrolytic environments.

### Cytocompatibility Testing

We evaluated the cytocompatibility of each hydrogel matrix by assessing the cell viability of both blank and MPs-containing gels. Hydrogel strips of standardized dimensions were punched from their respective monolithic slabs, and these strips were subsequently incubated with mouse 3T3 fibroblast cells at 37°C for 24 hours. It is worth noting that the size and dimensions of the hydrogel strips were intentionally maintained consistently with those used in our standard devices for *in vivo* insertion.

As depicted in **Figure 6**, both blank and MPs-containing alginate gel strips displayed minimal cytotoxicity towards the incubated fibroblast cells, with cell viability exceeding 90% for both gel types. A similar trend was observed for blank and MPs-containing GAC gels, where the cell viability for each gel type exceeded 80%.

**Figure 6:**
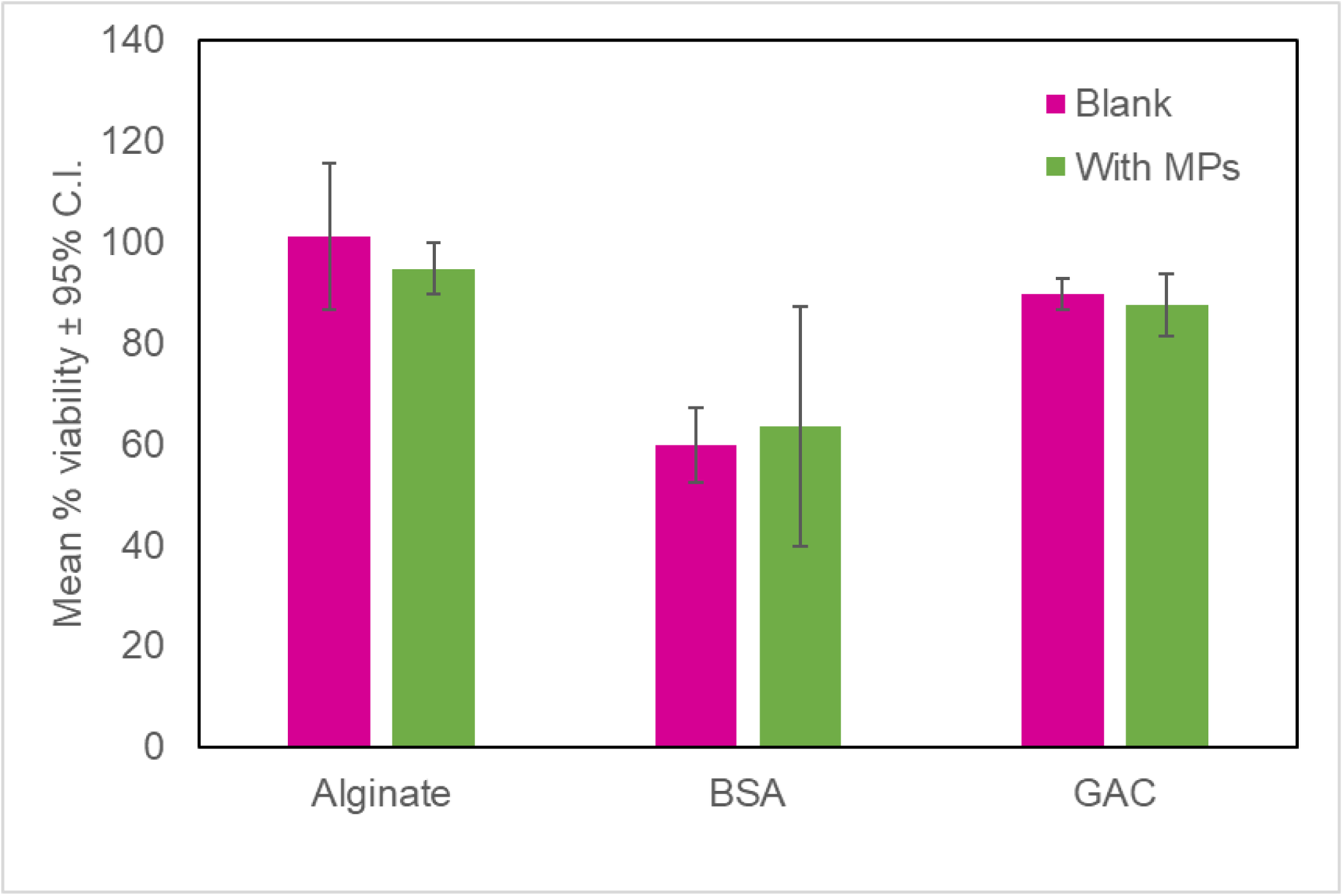
Cytotoxicity effects of blank and MPs-containing hydrogel strips [0.5 (w) X 0.5 (h) X 5 (l) mm] after 24 hours of incubation with 3T3 fibroblast cells. Both alginate and GAC samples (with and w/o. MPs) were least cytotoxic in nature, followed by BSA XLD samples.

However, the cell viability decreased to approximately 50% when exposed to blank BSA gels; this was further reduced to around 30% in the case of MPs-containing BSA gels. This complex behavior of BSA gels, with reduced cell viability, could be mainly attributed to the cytotoxic effects of unreacted glutaraldehyde present in the BSA gels [93] and, least likely, by antifouling properties of the BSA gels. [47], [94], [95], [96], [97] For future studies, such an issue can be mitigated by employing a pre-incubation step with BSA gels under a glycine-based inactivation buffer. [98] Alternatively, Ma and colleagues reported no such cytotoxicity issues with glutaraldehyde crosslinked BSA gels when sample gels were subject to a prior lyophilization cycle (to deactivate unreacted glutaraldehyde) before conducting cell culture studies. [49]

### Oxygen sensitivity studies of sensors in different natural hydrogel matrices

We conducted oxygen sensitivity studies to assess the quenching characteristics of phosphors embedded in the three natural hydrogel matrices. We incorporated 8.8 mg alginate microparticles (oxygen-sensitive) into three hydrogel matrices [1.5% calcium alginate, 20% BSA, gelatin-alginate-collagen (10-1-9%)] at a volume of 400 μL each. After crosslinking and subsequent gelation, we punched identical hydrogel strips (5×0.5×0.5 mm) from the slabs for sensor fabrication in quadruplicate. These sensors then underwent various sterilization and treatment procedures.

Air and nitrogen gas were mixed in a defined ratio to achieve varying dissolved oxygen (DO) levels between 0 and 257.9 µM in TRIS buffer (pH 7.2) at a constant temperature of 37°C. This range encompassed ∼17 µM of DO, which is representative of the oxygen concentrations found within human subcutaneous tissues. [99] Emission lifetimes were measured for each DO concentration, and the oxygen sensitivity was quantified by determining the Stern-Volmer constant (*Ksv*) in each case. The *Ksv* before and after treatment were compared, and the results are presented in **Figure 7**, with additional details provided in **Table 1**.

**Figure 7:**
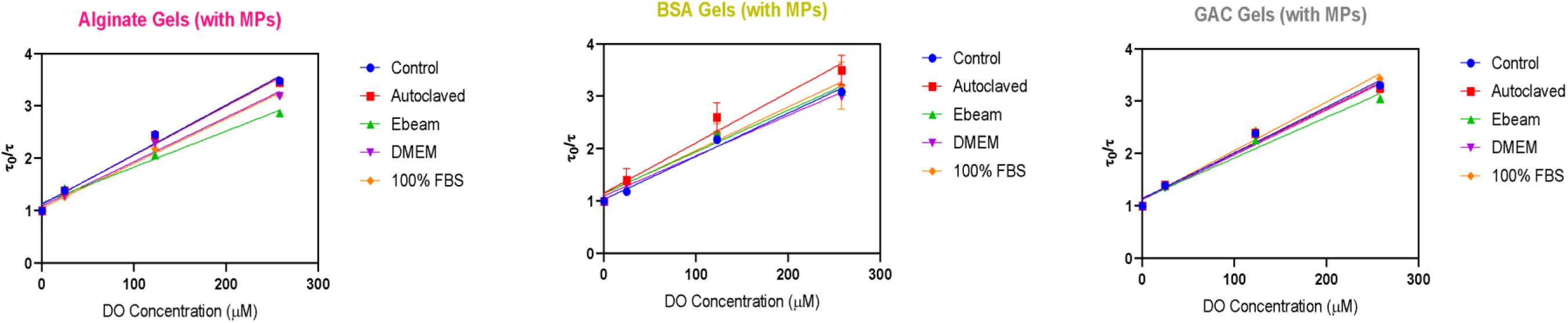
Stern-Volmer (SV) plot of oxygen sensors dispersed in alginate, BSA and GAC hydrogel matrices (n=3; 0.5 mm (w) X 0.5 mm (h) X 5 mm (l) strips) after exposure to different sterilization and treatment conditions.

**Table 1:**
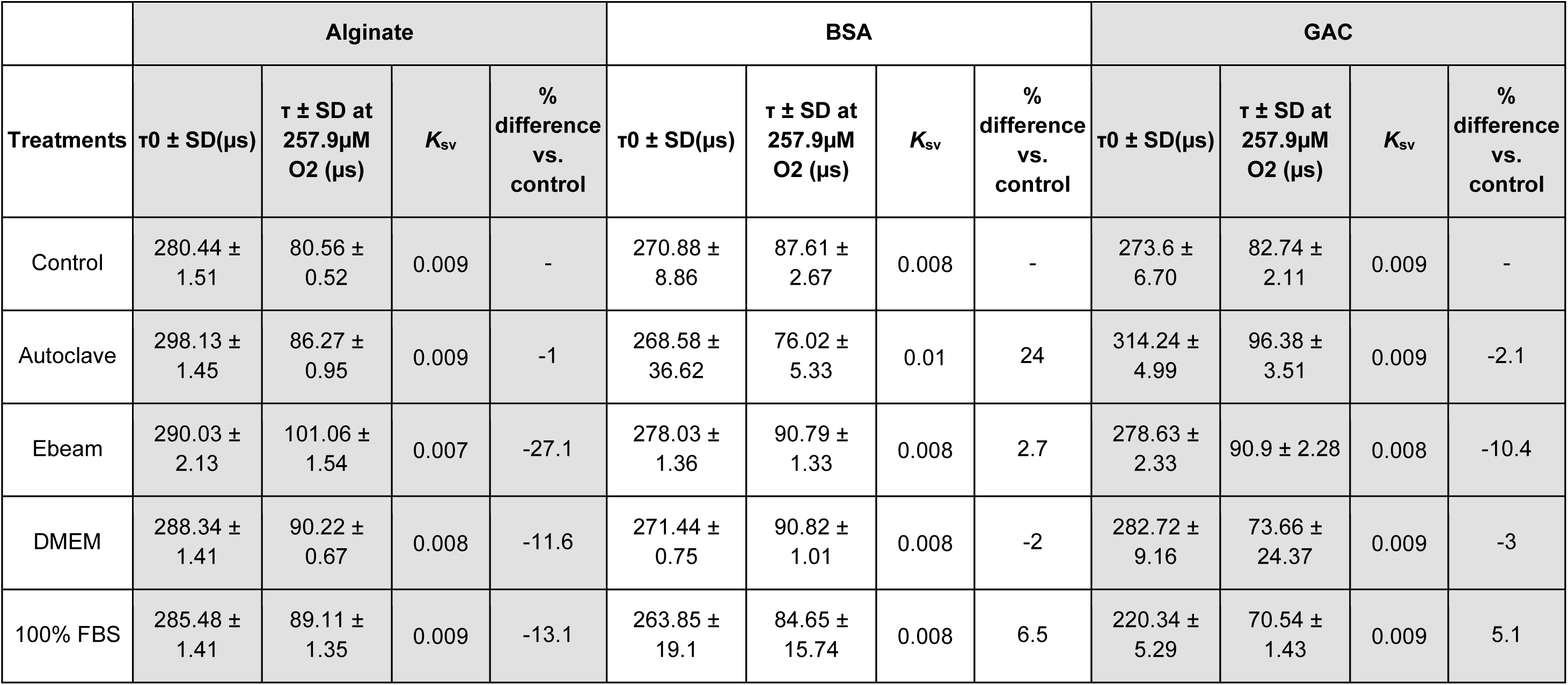
Oxygen sensitivity characterization and key Stern-Volmer testing parameters and lifetime values from all three sensors.

|  | Alginate |  |  |  | BSA |  |  |  | GAC |  |  |  |
| --- | --- | --- | --- | --- | --- | --- | --- | --- | --- | --- | --- | --- |
| Treatments | $\tau_0 \pm \text{SD}(\mu\text{s})$ | $\tau \pm \text{SD at } 257.9\mu\text{M O}_2 (\mu\text{s})$ | $K_{\text{SV}}$ | % difference vs. control | $\tau_0 \pm \text{SD}(\mu\text{s})$ | $\tau \pm \text{SD at } 257.9\mu\text{M O}_2 (\mu\text{s})$ | $K_{\text{SV}}$ | % difference vs. control | $\tau_0 \pm \text{SD}(\mu\text{s})$ | $\tau \pm \text{SD at } 257.9\mu\text{M O}_2 (\mu\text{s})$ | $K_{\text{SV}}$ | % difference vs. control |
| Control | $280.44 \pm 1.51$ | $80.56 \pm 0.52$ | 0.009 | - | $270.88 \pm 8.86$ | $87.61 \pm 2.67$ | 0.008 | - | $273.6 \pm 6.70$ | $82.74 \pm 2.11$ | 0.009 | - |
| Autoclave | $298.13 \pm 1.45$ | $86.27 \pm 0.95$ | 0.009 | -1 | $268.58 \pm 36.62$ | $76.02 \pm 5.33$ | 0.01 | 24 | $314.24 \pm 4.99$ | $96.38 \pm 3.51$ | 0.009 | -2.1 |
| Ebeam | $290.03 \pm 2.13$ | $101.06 \pm 1.54$ | 0.007 | -27.1 | $278.03 \pm 1.36$ | $90.79 \pm 1.33$ | 0.008 | 2.7 | $278.63 \pm 2.33$ | $90.9 \pm 2.28$ | 0.008 | -10.4 |
| DMEM | $288.34 \pm 1.41$ | $90.22 \pm 0.67$ | 0.008 | -11.6 | $271.44 \pm 0.75$ | $90.82 \pm 1.01$ | 0.008 | -2 | $282.72 \pm 9.16$ | $73.66 \pm 24.37$ | 0.009 | -3 |
| 100% FBS | $285.48 \pm 1.41$ | $89.11 \pm 1.35$ | 0.009 | -13.1 | $263.85 \pm 19.1$ | $84.65 \pm 15.74$ | 0.008 | 6.5 | $220.34 \pm 5.29$ | $70.54 \pm 1.43$ | 0.009 | 5.1 |

The Stern-Volmer plots depicted in **Figure 7** revealed no substantial differences in the oxygen sensitivity of MPs when dispersed in different matrices. Control gels for both alginate and GAC samples provided similar oxygen sensitivity responses with approximate Stern-Volmer quenching constant (K_sv_) values around 0.009 µM^-1^, while control BSA gels exhibited a comparable but slightly lower response as indicated by a K_sv_ value of 0.008 µM^-1^. Similarly, autoclaved alginate and GAC gels provided similar oxygen sensitivity responses with K_sv_ values around 0.009 µM^-1^, whereas autoclaved BSA gels reported significant differences in sensitivity with approximately 22% change against control BSA gels. It is also evident from the response of autoclaved BSA gels, especially under zero oxygen conditions, that these sensors exhibited the highest sensor-to-sensor variability (high SD values; see Table 1). We hypothesize that highly harsh physical conditions of autoclave (121℃ at 15 psi for 30 mins) could have resulted in complex changes in matrix structure, such as complete denaturation of BSA protein (at 90℃) [100] and accompanied by irreversible conformational changes in helices and aggregate formation. [101] Separately, we observed marked visual changes within post-autoclaved BSA gels, where the resulting gels were dark brown, shrank by 20-30%, and resulted in increased gel stiffness (data not shown). More investigation is required to understand better the underlying effects, which are beyond the scope of this study.

With electron-beam sterilized samples, significant differences in sensitivity were observed in oxygen sensors dispersed in alginate gels (∼24% change between controls and E-beam treatment) followed by GAC gels (∼12% change between control and E-beam treatment). No significant differences in oxygen sensitivity responses were observed with BSA gels. For other cell-culture related treatment conditions such as under incomplete cell culture medium (DMEM only) and 100% fetal-bovine-serum supplemented DMEM medium (100% FBS), only alginate gels provided significant difference in oxygen sensitivity responses with ∼12% change between control and DMEM treatment.

Based on the oxygen sensitivity profiles, BSA gels (except for autoclave response) and GAC (except for Ebeam) provided robust responses across various treatment methods. According to our findings, E-beam sterilization was the most appropriate method against BSA gels, whereas autoclave sterilization was used against GAC gels.

### Mechanical and Rheological Characterization Studies

#### 1. Mechanical Testing

Dynamic Mechanical Analysis (DMA) was used to assess the compression moduli of various hydrogel matrices in their blank and microparticle (MPs)-containing states, following specific treatment regimens. Quadruplicate samples were subjected to compression testing using a compression clamp, and statistical analysis was applied to evaluate compression modulus variations within each matrix category, as illustrated in **Figure 8**.

**Figure 8:**
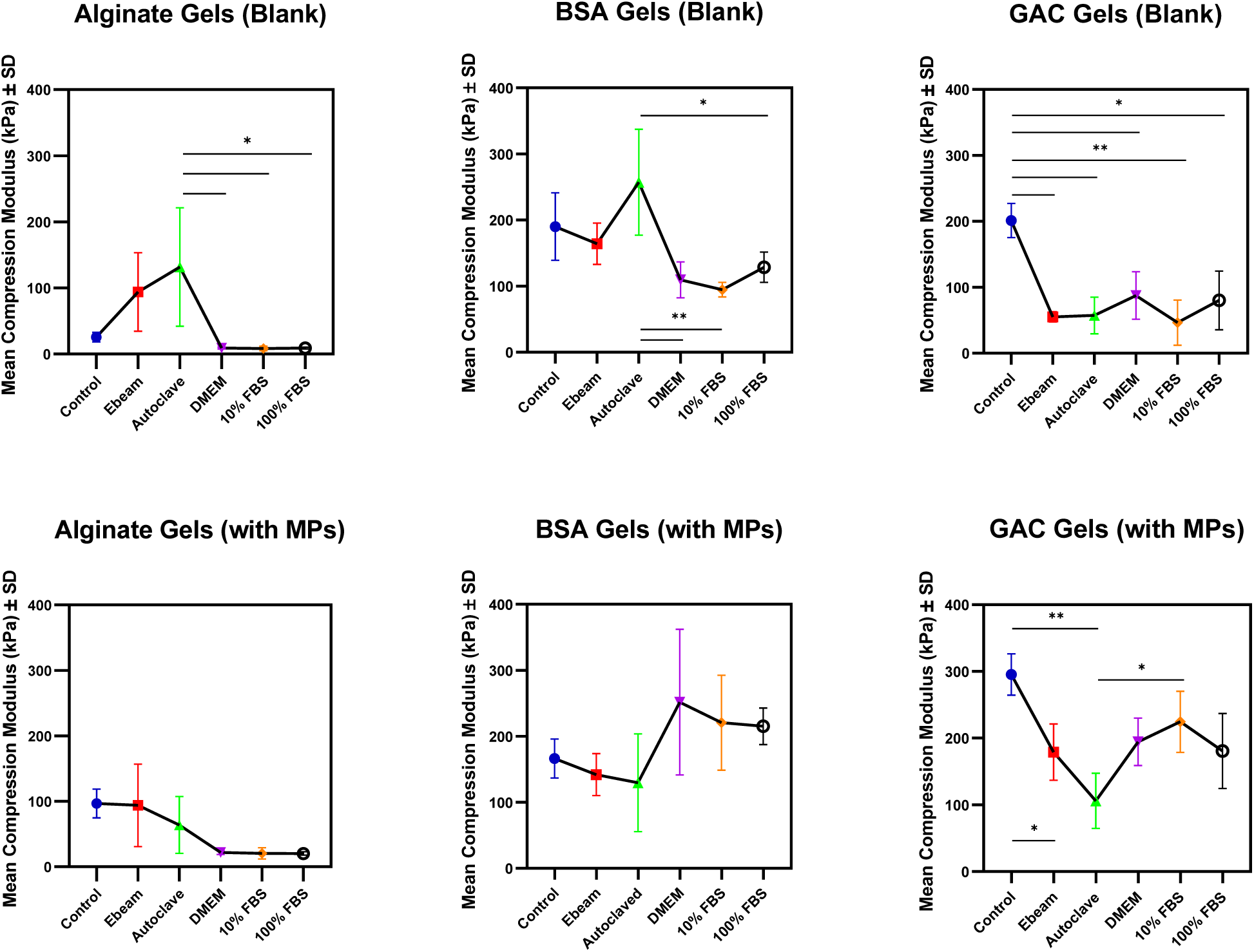
Comprehensive mechanical properties (compression modulus) and statistical analyses (1-way ANOVA) from N=3 samples from matrices under different treatment conditions

In the case of blank alginate and BSA gels, significant differences were noted among the treated samples, particularly when comparing autoclave treatment to DMEM, 10% FBS, or 100% FBS treatments. Furthermore, blank GAC gels exhibited higher variability among the treated samples, with significant differences evident in the mean compression modulus data, specifically between the control group and all other treatment conditions.

After incorporating MPs into the gels, a consistent trend was observed in alginate and GAC matrices. This trend indicated a relative increase in mechanical strength compared to their blank counterparts among the treated samples. Interestingly, for the BSA matrix, an increase in compression modulus was observed exclusively in MPs-containing gels subjected to DMEM, 10% FBS, and 100% FBS treatments. However, a slight reduction in the mean compression modulus was noted for control and Ebeam-treated samples. Notably, no statistically significant differences were discerned among the treated samples in alginate and BSA gels containing MPs.

In light of these mechanical profiles, we conclude that incorporating microparticles into these matrices has a minimal, if not reinforcing, effect on their mechanical strength.

#### 2. Rheological Testing

Rheological tests were conducted on hydrogel matrices comprising alginate, BSA, and GAC, encompassing both blank gels and those incorporating microparticles (MPs). These tests were administered after exposure to diverse treatment conditions, with the aim of elucidating alterations in their flow properties under varying testing conditions.

We initiated the experimentation with an oscillatory frequency sweep analysis, see **Supplemental Figure S1**. This procedure aimed to establish the maximum threshold frequency limit for each hydrogel material, whether in its blank or MPs-containing gels. This determination was crucial in assessing the material’s response to differing time scales, ensuring the complete restoration of samples to their original states before conducting subsequent rheological measurements. Subsequently, we conducted oscillatory strain sweep tests on each hydrogel material, encompassing blank and MPs-containing gels. The objective was to ascertain the threshold strain limits that would maintain the materials within linear viscoelastic (LVE) behavior, a fundamental aspect of their structural integrity. With the frequency and strain parameters established for each hydrogel matrix, we executed oscillatory time sweep tests to determine mean shear storage modulus (G’) and mean shear loss modulus (G") changes from triplicates, as plotted in **Figure 9 (a-b)**. In addition, **Figure 10 (a-c)** provides representative plots of storage vs. loss modulus changes over time from hydrogel samples following exposure to the specified treatment conditions.

**Figure 9a:**
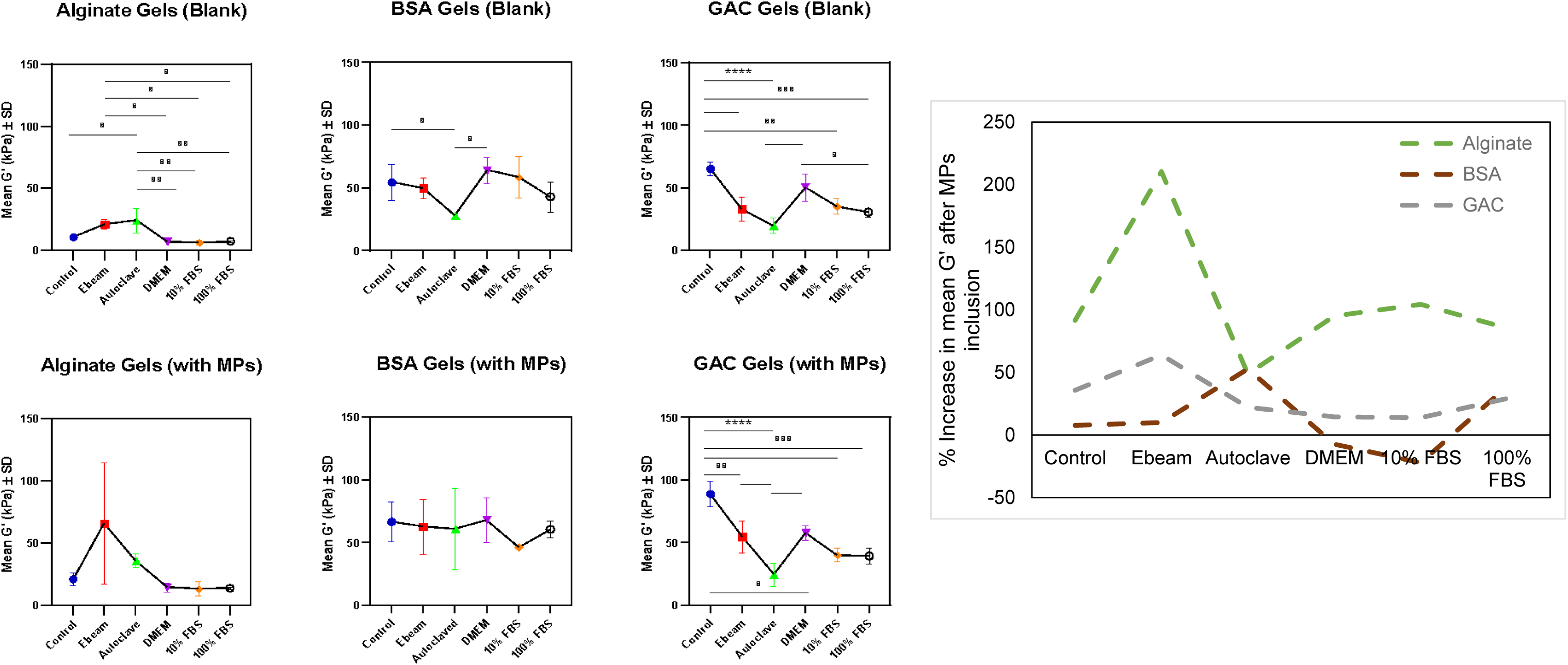
**(LEFT)** Storage Moduli (G’) changes and statistical analyses (1-way ANOVA) from N=3 samples of different matrices under different treatment conditions **(RIGHT)** Percentage increase in mean storage moduli of different matrices (with MPs) under different treatment conditions.

**Figure 9b:**
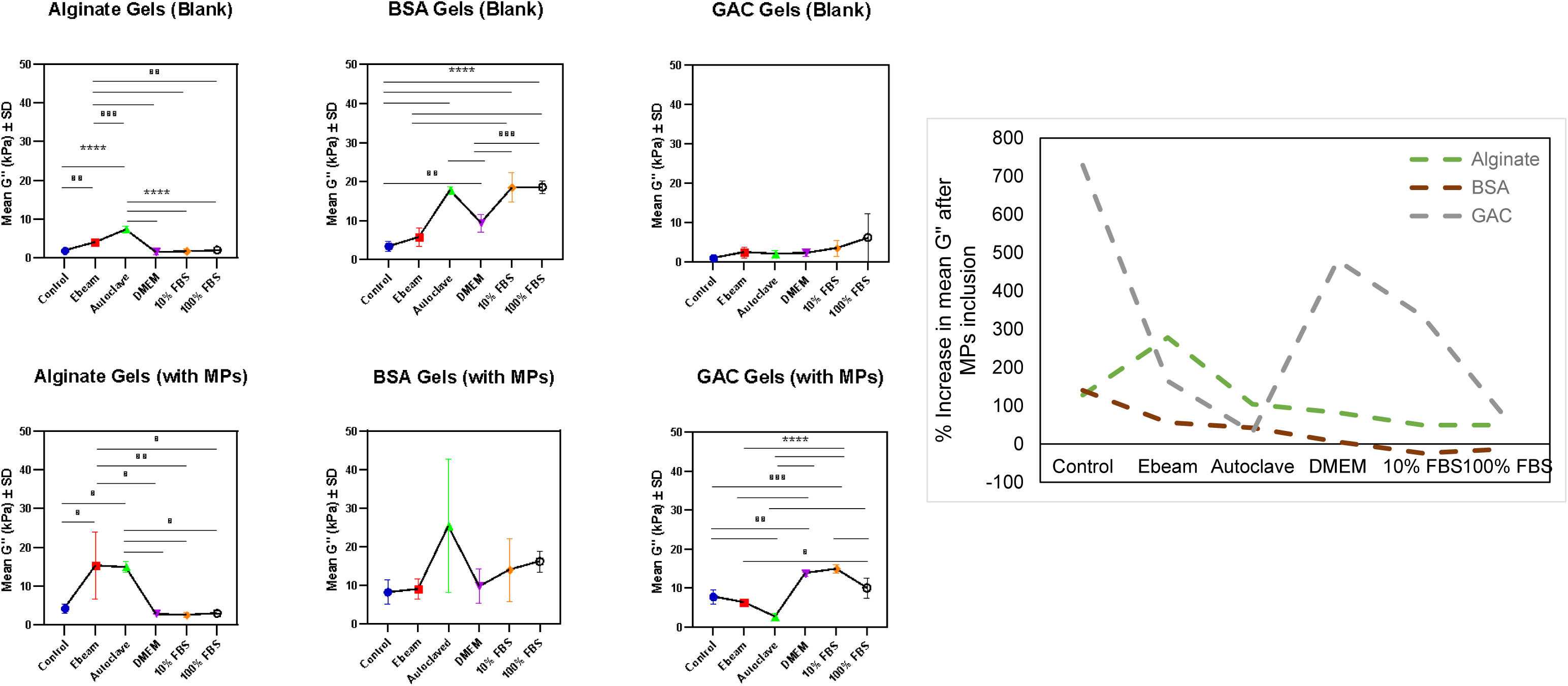
**(LEFT)** Loss moduli (G”) changes and statistical analyses (1-way ANOVA) from N=3 samples of different matrices under different treatment conditions **(RIGHT)** Percentage increase in mean loss moduli of different matrices (with MPs) under different treatment conditions.

**Figure 10a:**
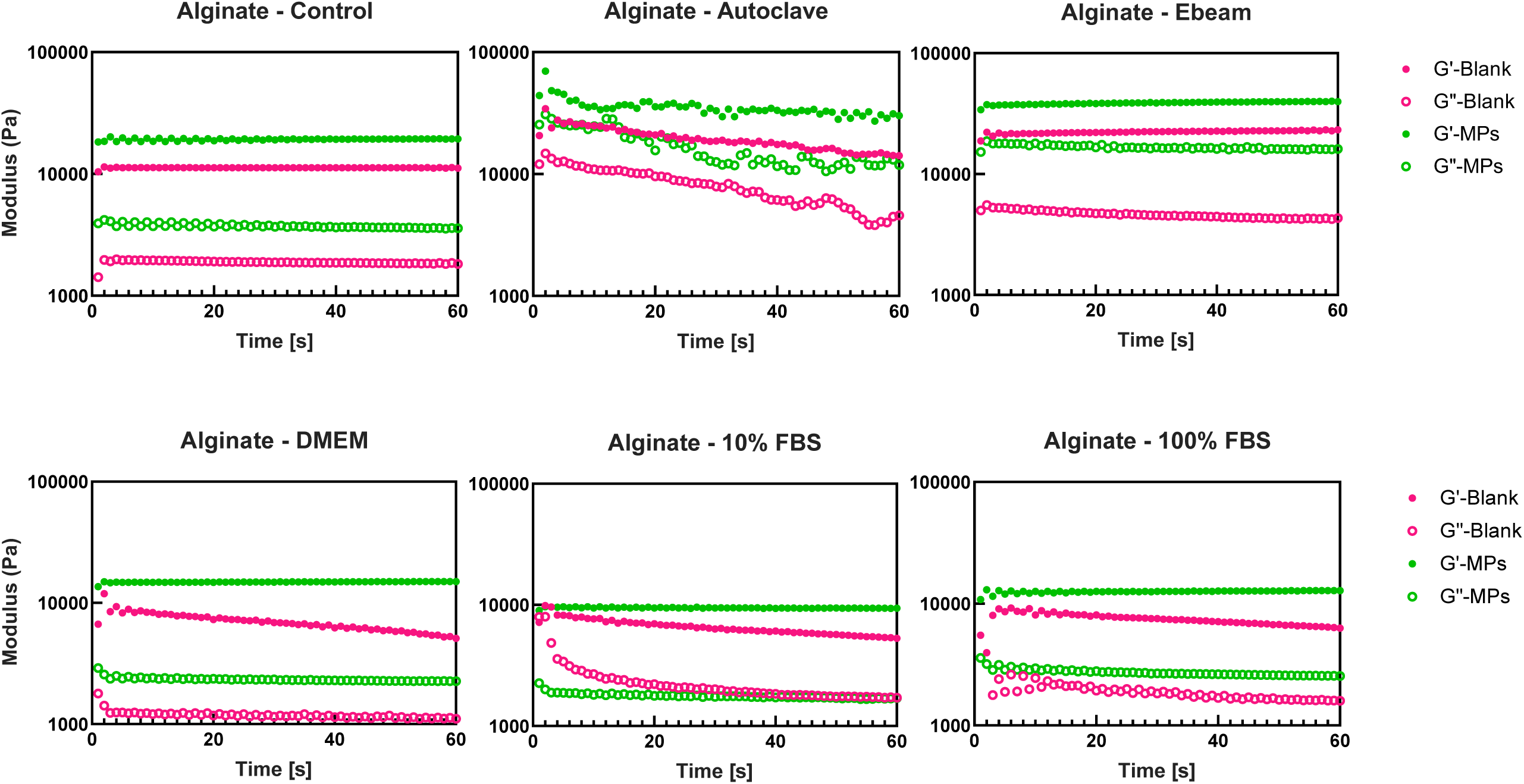
Representative oscillatory time sweep plots (G”/G” over time) from various treatment conditions – Alginate Gels

**Figure 10b:**
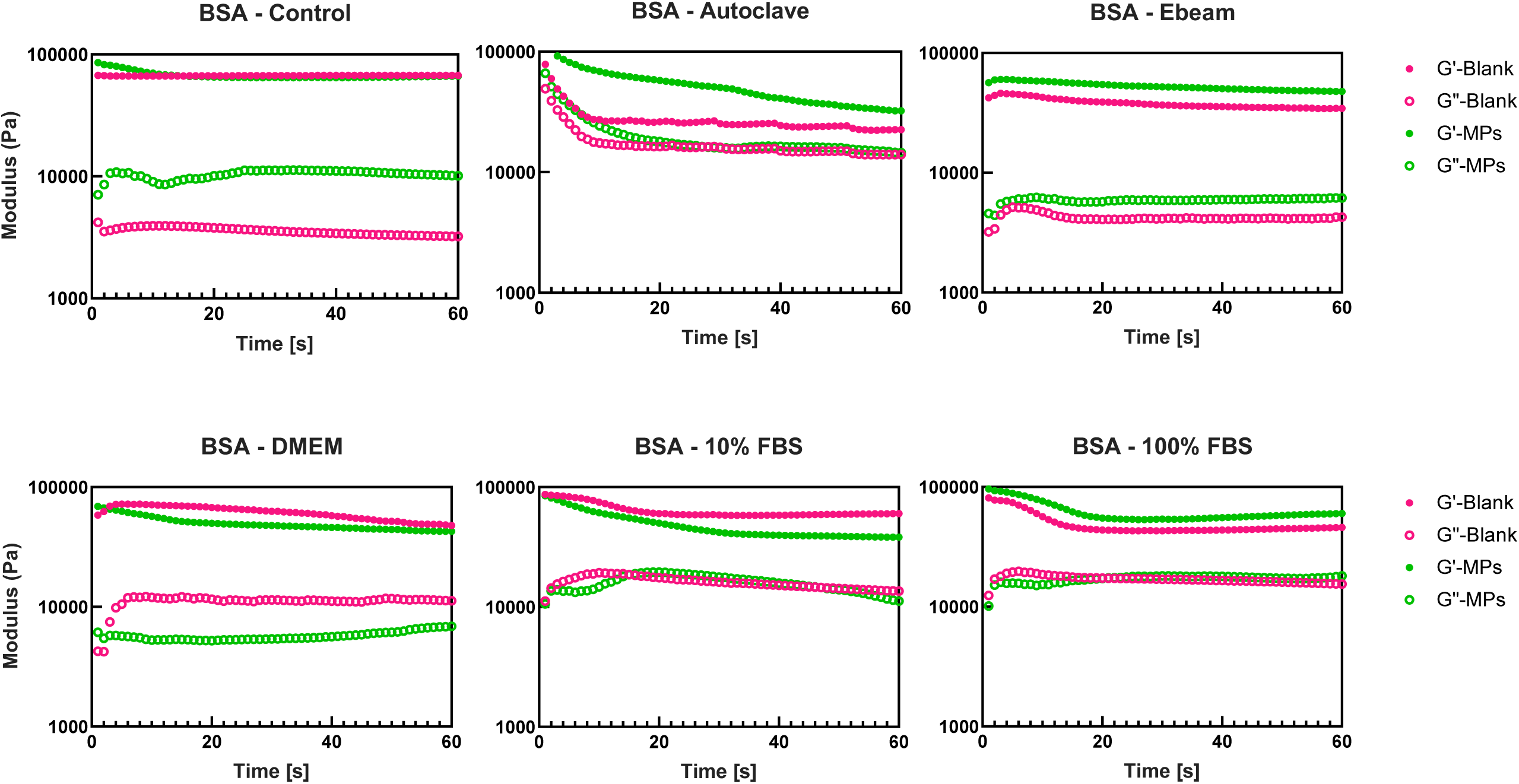
Representative oscillatory time sweep plots (G”/G” over time) from various treatment conditions – BSA Gels

**Figure 10c:**
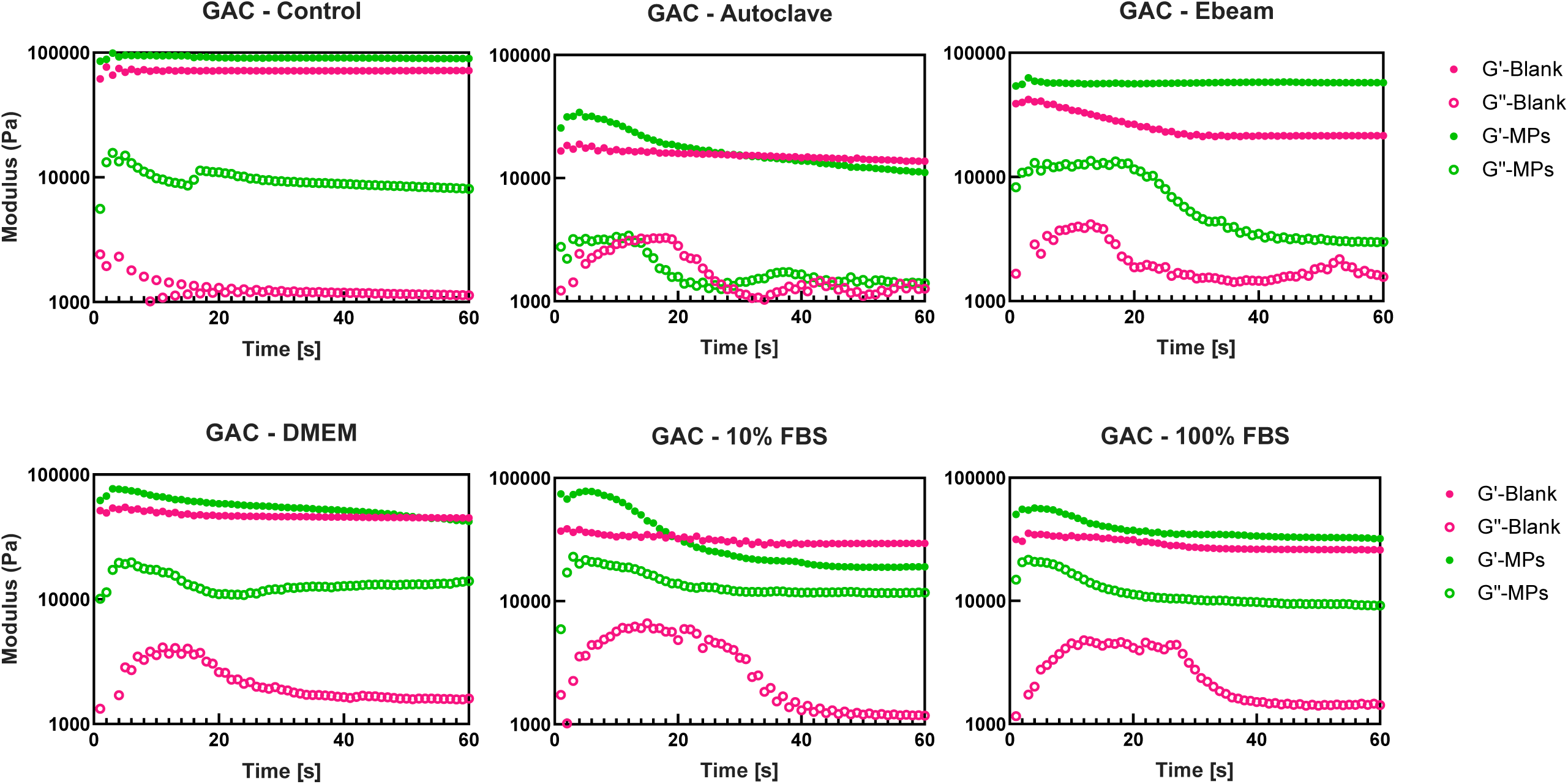
Representative oscillatory time sweep plots (G”/G” over time) from various treatment conditions – GAC Gels

Rheological assessments obtained through oscillatory time sweep analysis unveiled increased storage (G’) and loss (G") modulus properties. Both storage and loss moduli reflect material stiffness (elastic-like behavior) and energy dissipation (viscous-like behavior), respectively. As depicted in **Figure 9a**, except for BSA gels treated under cell culture conditions (DMEM and 10% FBS), the inclusion of oxygen-sensing alginate microparticles across all three natural hydrogels for all treatment conditions resulted in at least 7% (BSA gels) to >90% (alginate gels) increase in G’ values against the blank gels. These trends were also evident through our DMA analysis (see Mechanical Testing Section), thus validating our observations that including microparticles does lead to strengthening alginate and GAC gels. Some logical explanations for this behavior can be 1) additional crosslinking sites imparted by microparticles, leading to a stiffer and denser network, and 2) adding rigidity and hindering deformation through physical entrapment of microparticles within the hydrogel network. [90], [102]Regarding different treatment conditions, MPs-containing alginate and BSA gels showed no statistically significant difference in their mean *G’* values following various treatment conditions. Notably, for MPs-containing GAC gels, statistically significant differences in *G’* were observed between control against all treatment conditions. Similarly, blank GAC gels also reported statistically significant differences in G’ changes between control against all treatment conditions (except DMEM). For blank alginate gels, statistically significant G’ changes were observed between control vs. autoclave; Ebeam vs. DMEM, 10% FBS and 100% FBS; and autoclave vs. DMEM, 10% FBS and 100% FBS conditions were observed. Only blank BSA gels provided the most consistent G’ properties, with significant differences only observed between control vs. autoclave and autoclave against DMEM treatment conditions. As discussed earlier, the autoclave sterilization method is expected to result in irreversible conformational changes in BSA gels. Therefore, except for autoclave-treated BSA gels, we conclude that both blank and MPs-containing BSA gels provide the most consistent rheological behavior.

**Figure 9b** represents a comparison of loss modulus (G") analyses between blank and MPs-containing natural hydrogels following different sterilization and treatment conditions. G" changes provide insight into possible energy dissipation mechanisms, where higher G" is indicative of higher energy dissipation as heat and more viscous-like gel behavior. Firstly, the inclusion of microparticles in all hydrogel matrices, excluding BSA gels under DMEM, 10% FBS, and 100% FBS, resulted in at least a 100% (alginate and BSA gels) to 700% (GAC gels) increase in G" values. One possible explanation for such increased viscous-like behavior is higher perforations in alginate and GAC hydrogels (see SEM data) and higher swelling ratios against BSA gels, facilitating free solvent movement across the hydrogel mesh and effective energy dissipation. [103] Secondly, upon individual samples were compared against different treatment conditions, such as for both blank and MPs-containing alginate gels, a significant difference in G" was observed among control vs. Ebeam, autoclave; Ebeam vs. DMEM, 10% FBS and 100% FBS; and autoclave vs. DMEM, 10% FBS and 100% FBS conditions. For blank BSA gels, significant differences were observed among control vs. autoclave, DMEM, 10% FBS, and 100% FBS conditions; Ebeam vs. 10% FBS, 100% FBS; autoclave vs. DMEM; and DMEM vs. 10% FBS and 100% FBS conditions. No significant differences in G" changes were observed in MPs-containing BSA and blank GAC gels when exposed to various sterilization and treatment conditions. Finally, for MPs-containing GAC gels, significant differences in G" were observed between control vs. autoclave, DMEM, 10% FBS; Ebeam vs. DMEM, 10% FBS, 100% FBS; autoclave vs. DMEM, 10% FBS, 100% FBS; and 10% FBS vs. 100% FBS conditions. As mentioned earlier, higher variabilities in G’ and G" changes within alginate and GAC gels could be explained by various involved mechanisms in such gels, such as physical disruption of the continuous polymeric network [89], [90], swelling ratio difference between microparticles/bulk gel skewing overall matrix concentration [91], and matrix compositional changes affecting crosslinking density [92].

Another perspective for considering our MP-containing gels’ stability and mechanical behavior would be to consider them hydrogel microparticles (HMP) composites. [90] HMPs are fabricated similarly, where microparticles are incorporated into a secondary hydrogel. Unless there is a substantially higher concentration of embedded microparticles, the mechanical properties of secondary hydrogels dominate the composite system’s behavior. Researchers have exploited these materials to engineer mechanically tough hydrogels by either preventing crack propagations in the material by blunting it with microgels [104] or by incorporating stiffer microgels in otherwise soft hydrogels to increase its toughness. [105]

Consistent with our findings from compression testing, incorporating microparticles into all three matrix types yielded a noticeable increase in storage and loss moduli, with blank and MPs-containing BSA gels providing the most consistent rheological behavior.

## Conclusion

We explored diverse natural hydrogels to design a conformal, sterilizable, and functionally stable insertable optical oxygen biosensor. Based on oxygen sensitivity profiles, E-beam sterilization proved most effective against BSA gels, while autoclave sterilization was an effective choice against GAC gels. In terms of mechanical properties, the compression modulus of our oxygen sensors ranged from approximately 0.017 MPa (alginate with MPs) to about 0.33 MPa (GAC with MPs). Variations in hydrogel properties were expected due to the inherent variability of natural biomaterials and the sensitivity of the crosslinking process. Interestingly, microparticle inclusion within natural hydrogels significantly increased alginate and GAC gels’ mechanical strength and energy dissipation mechanism. Notably, BSA gels exhibited the most consistent mechanical and rheological properties.

Based on **Figure 1**, studies revealed significant variability in whole-skin elastic modulus (E) across different body regions, age groups, and sexes. Moreover, the same anatomical site can exhibit substantial elastic modulus value variations due to differences in measurement techniques. Given the diverse anatomical, physiological, and biomechanical considerations, a rigid "one-size-fits-all" solution cannot effectively address the optimal insertable biosensor design challenge. Therefore, we propose exploiting the inherent variability in the elastic modulus of skin against insertable biosensors, selecting sensors specifically suited for optimal tissue compatibility, thereby enhancing their long-term performance and reducing the risk of rejection. The wide range of material choices and tunable mechanical properties empower researchers to design tissue-compatible devices. These features hold significant promise for long-term applications.

### Experimental Section Methods and Materials

**Figure 2** provides a schematic representation of the experimental layout for this study. Sodium alginate (Cat No. A215875000 - MW: 100000 Da), calcium chloride (Cat No. 22350), calcium carbonate (Cat. No. 239216), TRIS base (Cat No. T1503), and catalase (CAT) from bovine liver (Cat. No.C9322), and alginic acid sodium salt, glutaraldehyde (25% in water), and glycine were purchased from Sigma-Aldrich, Inc., St. Louis, MO. Collagen (rat tail-type I) and neutralization solutions were purchased from Advanced Biomatrix, Carlsbad, CA. Bovine serum albumin, BSA (Fraction V, Protease-Free), was purchased from GoldBio, St Louis, MO. Trizma hydrochloride (TRIS HCl) (Cat No. VWRB85827), BIS acrylamide (Cat No. 0172, Amresco), Gelatin (Pork Skin Type A), N-hydroxysuccinimide (NHS) and 1-Ethyl-3-(3-dimethyl aminopropyl) carbodiimide (EDC), and phosphate buffer saline (PBS) were purchased from VWR Chemicals LLC, NY, USA. 2-(methacryloxy) ethyl phosphorylcholine MPC (Cat. No. M2005), glucose oxidase (GOx) from *Aspergillus niger* (Cat No. G0050)), (Z, Z, Z)-Sorbitan tri-9-octadecenoate (SPAN 85) (Cat. No. S0064), Polyoxyethylenesorbitan Trioleate (TWEEN 85) (Cat. No. T0547) were purchased from Tokyo Chemical Company Co., LTD, Tokyo, Japan. Warrington, PA. Palladium (II) meso-tetra-(sulfophenyl) tetrabenzoporphyrin sodium salt (Cat No. T41161 - MW: 1327.55 g/mol) was purchased from Frontier Specialty Chemicals, Logan, UT. Isooctane (Cat No. 94701) was purchased from Avantor Performance Materials, LLC, Randor, PA. All the above chemicals were used as obtained without any further purifications. Dulbecco’s Modification of Eagle’s Medium (DMEM) with L-glutamine, 1g/L glucose, and sodium pyruvate) was obtained from Corning/Mediatech, Inc., Manassas, VA, USA. Fetal Bovine Serum (FBS; heat activated) was procured from Gibco, Grand Island, NY, USA. EmbryoMax® penicillin/streptomycin (P/S) antibiotics and cell counting kit (CCK-8; Cat. No. 96992) for cytotoxicity assay were purchased from Millipore Sigma St. Louis, MO, USA.

### Preparation of oxygen-sensitive alginate microparticles

Oxygen-sensitive alginate microparticles were fabricated using the water-in-oil emulsification method, as described previously. Briefly, 5 ml of 3% w/v alginate (in DI water) solution was prepared and mixed overnight under nutation with 500 μL of 10 mM phosphor dye (in DMSO). Next day, in a 50 ml centrifuge tube, add 260 μL of SPAN 85 surfactant into 10.8 mL isooctane solution and mix thoroughly by vortexing the mixture. In a 2 mL centrifuge tube, 130 μL of TWEEN 85 surfactant was added to 1.5 mL of isooctane solution. Similarly, 4 ml of 10% w/v calcium chloride (in DI water) solution was prepared. The homogenizer speed was set at 8000 rpm, and SPAN 85/isooctane solution was spun for 10 seconds, and then the alginate-phosphor dye mixture was slowly added. After 10 seconds of homogenization, the TWEEN 85/isooctane mixture was added and homogenized for 10 seconds, followed by CaCl_2_ solution, and homogenized for an additional 15 sec. Finally, the resulting emulsion was transferred to a round bottom flask with a magnetic stir bar and stirred for 20 mins at 300 rpm at room temperature. The emulsion was transferred to a 50 mL centrifuge tube and centrifuged at 625 *g* for 2 mins; the supernatant was discarded, and to the pallet, 1 mL of DI was added to resuspend the particles. Next, the solution was re-centrifuge at 625 *g* for 1 minute, rewashed with DI H2O, and resuspended in 1 mL PSS Wash solution. At each washing step, pallets were thoroughly dispersed using pipette tips - breaking the aggregate formation. Next, the solution was transferred to 2 mL centrifuge tubes, and the layer-by-layer (LbL) step was proceeded with.

### The layer-by-layer (LbL) nanofilm coating procedure

LbL solutions and procedures were applied using a previously established procedure[8]. Microparticles under PSS wash solution were briefly centrifuged, the supernatant was discarded, and 1 mL of PAH solution was added. The pellet was dislodged using a pipette tip by trituration of solution 5-6 times, followed by centrifugation. The supernatant was discarded, and the pellet was resuspended in 1mL of PAH Wash solution. Following the same trituration method, the solution was centrifuged and resuspended into 1 mL of PSS solution, triturated, and then re-centrifuged, followed by resuspension and trituration into 1mL of PSS wash buffer. This completes the first bilayer step. The process was repeated until five bilayers were achieved, with the final PSS Wash step repeated twice. The dry weight of the microparticle suspension was determined by taking the empty weight of centrifuge tubes (in triplicates), then adding 30uL each of the microparticle suspension into each tube and drying it overnight under the vacuum oven (at 60℃). The stock microparticle suspension was stored under the dark in the PSS Wash buffer at 4℃. The next day, the dried weight of each sample was calculated by subtracting the dried tube weight from the empty tube weight.

### Size Distribution Analysis

Cellometer was used to perform microscopy of alginate microparticles and to determine the size distribution. First, stock microparticle solution was diluted 1:25 in TRIS buffer (pH 7.2) with 10mM CaCl_2_ solution, and then 20 µL volume was injected into the slide.

### Degradation Studies

For degradation studies, blank hydrogel samples (6 mm dia. X 0.75 mm thickness), in triplicates, were soaked in 5ml of 100 mM Phosphate Buffer Saline (PBS) solution for three months. Images were taken on day 0, day 15, one month, and three months, respectively.

Accelerated chemical degradation studies were adapted from Briggs et al.[106], where blank and MPs-containing hydrogel punches [3 (*dia.*) x 0.75 (*h*) mm] in triplicates from all three matrices were individually weighed (hydrated) on day 0 before incubated under 1 mL of 0.1 M sodium hydroxide solution. The solution was decanted on day 5, and gels were weighed (hydrated) and re-incubated in fresh NaOH solution until the next time point.

### Preparation of calcium alginate hydrogel slab with oxygen-sensitive MPs

For the preparation of calcium alginate hydrogel, 75 µL of microparticle suspension (8.8 mg of MPs) was mixed with 25 µL of 33.3 % aqueous suspension of calcium carbonate and 200 µL of 3.0 % aqueous solution of sodium alginate. Finally, 100 µL of MES buffer with pH 6.1 was added, and the whole mixture was quickly transferred into a rectangular mold made with microscope slides and a 0.75 mm Teflon spacer. The calcium release from the carbonate induced gelation of the alginate, which was allowed to proceed for 20 minutes. The resulting oxygen-sensing alginate hydrogel slab was carefully recovered and stored for 24 h in the TRIS buffer (pH 7.2) with 10.0 mM calcium chloride at 4°C before testing.

### Preparation of Gelatin-Alginate-Collagen (GAC) hydrogels with MPs

Two separate solutions, 25% gelatin and 5.25% alginate, each in 10mM MES buffer (pH 6.1), were prepared by heating and stirring the solution to 60°C for 5 mins. Solutions were cooled down under the water bath and held at 37°C. A solid mass of NHS was added to the alginate solution at 5.23 mg/ml concentration and dissolved using a stir bar. Next, to 75 uL of microparticle suspension (8.8 mg dry weight), 160 µL of gelatin and 76.2 µL of alginate solutions were added, followed by 88.9uL of neutralized collagen solution (0.82 mg/ml final concentration). The mixture was briefly vortexed and quickly transferred to 37°C in a water bath. The mixture was collected and injected between two glass slides sandwiched by a 0.75 mm Teflon spacer. Slides were cooled to 4°C for 10 mins and then submerged in a Petri dish pre-filled with EDC solution (10mg/ml) prepared using 10mM MES buffer (pH 6.1). The dish was covered with light and stirred for 2 hours. Finally, the gel was washed twice with DI water, covered, and stored in PSS Wash buffer at 4°C until the subsequent use.

### Preparation of Bovine Serum Albumin (BSA) hydrogels with MPs

A 30% stock solution of BSA in 100 mM PBS buffer (pH 7.4) was prepared and stored at 4°C unless used. To 266.7 µL of 30% BSA solution, 75 uL of microparticle suspension (8.8 mg dry weight) and an additional 58.3µL of PBS buffer were added to achieve the final concentration of BSA at 20%. The mixture was nutated for 15 mins, followed by the addition of 16 µL of glutaraldehyde stock solution, pipette mixed for 1-2 times and quickly filled between two glass slides (paraffin wrapped) sandwiched with a 0.75mm Teflon spacer. The polymeric gel was left to crosslink under room temperature for 15 mins. Finally, the gel was washed twice, once under 10ml of 1% glycine solution followed by PSS WASH buffer, covered and stored in PSS Wash buffer at 4°C until the subsequent use.

### Characterization of oxygen sensitivity

Oxygen sensitivity was determined by measuring phosphorescence lifetime under exposure to different dissolved oxygen concentrations. Circular discs (3 x 0.75 mm, n = 4) from each MP-containing hydrogel matrix were loaded inside a rectangular flow cell with one side dedicated to inlet and outlet openings for buffer tube connections, while the other side has windows dedicated to the attachment of optical readers for readouts. Hydrogel samples were loaded inside the flow cell using an acrylic sheet that holds the samples using rubber cantilevers and then fixed and sealed inside the flow cell where they can be optically interrogated from outside through windows by placing optical readers next to the acrylic/window interface. A pair of digitally controlled mass flow controllers (MKS instruments PR4000B controller) was used to mix externally supplied air and nitrogen flow in a defined ratio between 0 and 21% for the final achievement of the dissolved oxygen (DO) levels between 0 and 257.9 µM. The pre-mixed air-nitrogen gas was first flushed and mixed in a round bottom flask filled with TRIS buffer solution, followed by buffer circulation towards the samples in the flow cell using a recirculating flow system described previously. [107] Finally, the entire flow cell and buffer solution assembly were placed inside an incubator at 37°C.

As the oxygen concentration varied, an established time-domain phosphorescence lifetime reader system (ex: 630 nm, em: 800 nm) was used to record the lifetime from each hydrogel sample at 10 sec of intervals. Later, the Stern-Volmer constants were calculated using the following equation (1):

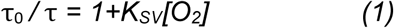

Where τ_0_ and τ represent phosphorescence lifetimes under the absence and presence of oxygen, respectively, at a given oxygen concentration [O_2_], and the Stern-Volmer constant (K_SV_) represents the slope of the linear relationship between the ratio *τ*_0_/*τ* and [O_2_]. K_SV_ values were extracted by drawing the best-fit line after plotting the phosphorescence lifetime ratios against varying oxygen concentrations.

### Cytotoxicity studies of blank and oxygen-sensitive (MPs containing) hydrogels

NIH/3T3 (mouse embryonic fibroblasts) cells at passage 24 were generously gifted by Dr. Daniel Alge from Texas A&M University. Cells were grown in DMEM culture media supplemented with 10% FBS and penicillin/streptomycin antibiotics (1X) and maintained in a humidified incubator with a 5% CO_2_ supply at 37°C. First, a cytotoxicity assay with blank and oxygen-sensitive hydrogel strips was performed according to the manufacturer’s protocol. After achieving 80% cell confluency, the 3T3 fibroblasts were trypsinized and seeded into the 96 well-plates at a cell density of 5000 cells/well in 100µL of DMEM media for 24 hours. Next day, hydrogel strips (0.5 × 0.5 × 5 mm) from blank and MPs gels (O_2_-sensitive) were punched in triplicates, soaked in sterile PBS solution in Petri dishes, and exposed to UV light of biosafety cabinet for 20 mins. Strips were later transferred to designated wells of the 96 well-plate (with cells) and incubated for 24 hours. The following day, 10uL of CCK-8 reagent was added to each well, followed by 4 hours of incubation, and absorbance measurement at 450 nm was recorded.

### Scanning Electron Microscopy

JEOL JCM-5000 Neoscope under high vacuum at 10 kV was used to image the surface topology of various hydrogel samples. Before imaging, hydrogel samples were freeze-dried for three days to remove moisture content.

### Swelling Properties and % EWC determination

Circular discs (6 × 0.75 mm) from each hydrogel sample (in triplicates) were punched out. After weighing out blank tubes, the weight of swollen hydrogels (*W_s_*) was recorded within individual tubes. Samples were freeze-dried for seven days, followed by measured dried weights (*W_d_*). The swelling ratio was calculated using the given formula:

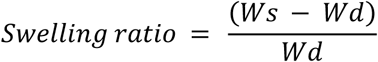

Hydrogel samples were reswelled in 1mL of TRIS buffer (7.2) with 10mM CaCl_2_ overnight. The next day, an excess buffer from each tube was removed, gels were gently damped with Kim wipe paper at their edges, and weights were recorded to calculate the re-swelled ratio and % EWC of samples before and after one freeze-drying cycle.

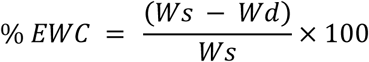

### Compression Testing

Compression moduli were determined through Dynamic Mechanical Analysis (DMA) using a TA Q800 instrument with compression clamps. Circular discs [n=4; 8 (dia) X 1.5 (h) mm] from both blank and MPs-containing hydrogels were punched out. Multi-stress/strain test was performed where samples were constantly held at room temperature and frequency of 1 Hz, while strain was varied from 0.0001 to 1% (amplitude from 0.1 to 15 μm). These parameters were critical to maintaining the linear viscoelastic region (LVE) within sample materials while measurements were carried out.

### Rheological Testing

Oscillatory frequency sweep, strain sweep, and time sweep experiments under controlled strain deformation mode were performed using an Anton Parr Rheometer with PP08 disposable plates, and shear storage and shear loss moduli were determined. Samples from both blank and MPs-containing hydrogels were punched into circular discs [n=4; 8 (dia) X 1.5 (h) mm]. All tests were performed at room temperature, with a force of 0.5 N. For each hydrogel type with oscillatory frequency sweep experiments, 1% strain was applied, and frequencies were ramped between 0.1 and 10 Hz to determine the maximum threshold frequency limits. The dedicated frequency limit against each hydrogel type (1 Hz for BSA and 0.5 Hz for alginate and GAC) was used to carry out an oscillatory strain sweep by ramping strains between 0.1 and 10%. Similarly, the threshold limit for strain % was determined against each hydrogel type to remain within the LVE regions. An oscillatory time sweep (1 min) was carried out at 1% strain at 1 Hz for BSA, 0.1% strain and 0.5 Hz for alginate, and 0.5% strain and 0.5 Hz for GAC, respectively.

## Acknowledgments

Some of the images in the article were created with Biorender. Research reported in this paper was supported by National Institutes of Health under award number R01EB024601

---

**Supplemental Figure S1:**
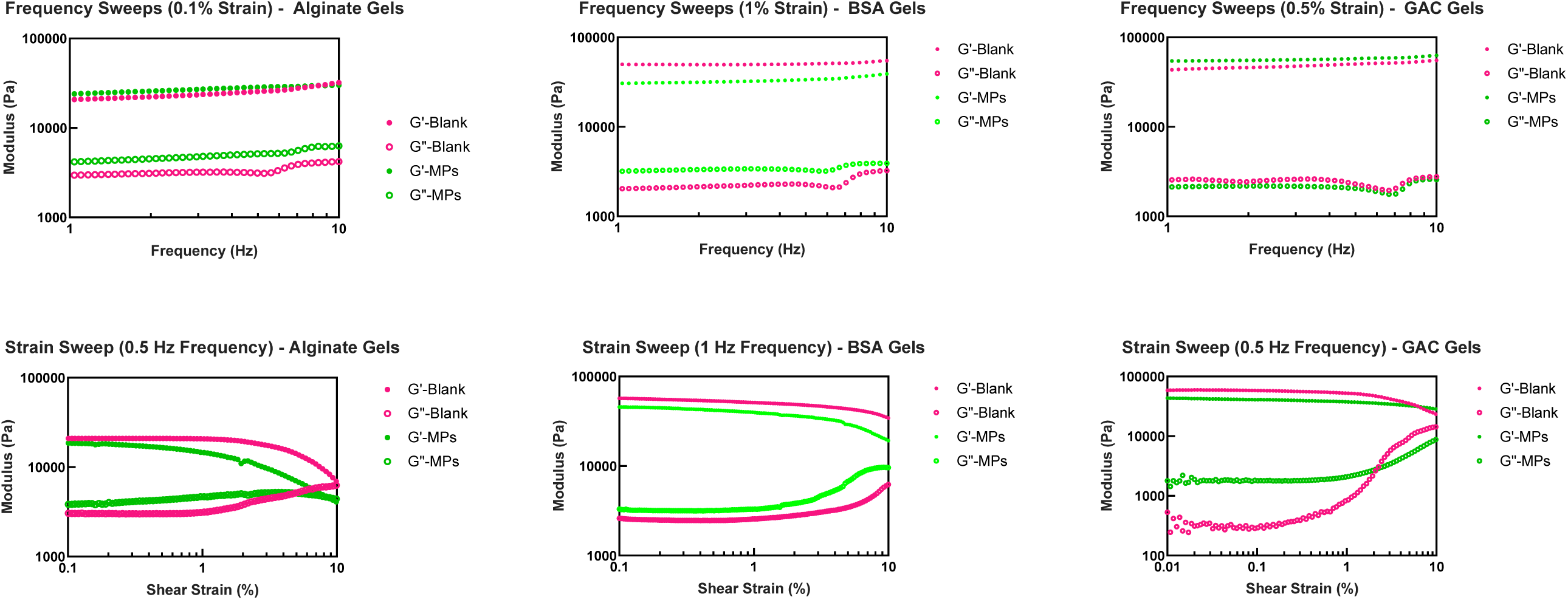
Representative frequency and strain sweeps plots from all three matrices

## Notes

### Competing Interest Statement

The authors have declared no competing interest.

